# Genetic background influences the effects of RNAi-by-feeding for DNA methyltransferases in *Daphnia pulex*

**DOI:** 10.1101/2025.10.02.680119

**Authors:** Trenton C. Agrelius, Krista Harmon, Allison Reed, Samuel B. Burnett, Rekha C. Patel, Jeffry L. Dudycha

## Abstract

DNA methyltransferases (DNMTs) are well-characterized epigenetic enzymes responsible for transferring methyl groups to and from DNA. Three main DNMT orthologues differ in function and methylation capability. They each are evolutionarily conserved across diverse taxa, but few studies investigate them jointly. We did so in *Daphnia*, freshwater microcrustaceans used extensively in research on genetic diversity, phenotypic plasticity, and maternal effects. Most *Daphnia* reproduce asexually through cyclic parthenogenesis, making the *Daphnia* system an ideal choice for studying epigenetic phenomena. Advances in gene expression control techniques, including RNA interference (RNAi), have increased the versatility and power of the *Daphnia* system. RNAi is a post-transcriptional gene silencing mechanism that operates through sequence-specific cleavage of endogenous messenger RNA (mRNA) transcripts. Here, we used an RNAi bacterial feeding regime to target the three DNA methyltransferase genes in two clones of *Daphnia pulex.* We observed significant genotypic differences in response to the RNAi feeding regime, namely, the mortality of one clone. In the other, DNMT expression significantly increased in five of the six experimental treatments, with the highest level observed in animals treated with the GFP double-stranded RNA bacterial vector control. However, DNMT expression was reduced in all three DNMT RNAi treatments relative to the GFP control. Furthermore, we found strong cross-reactivity, where targeting one DNMT resulted in the reduction of the other two. This response may be associated with known immune pathways involving signal transduction that can be stimulated by viral and bacterial signals, or it may result from previously unknown aspects of DNMT biology.

## Introduction

DNA methyltransferases (Dnmts) are epigenetic enzymes that add methyl groups (-CH3) to the 5’-carbon in cytosines (Goll and Bestor 2005). There are at least three well-characterized orthologues that differ in function and methylation capability. DNMT1 is a highly conserved methyltransferase gene that maintains DNA methylation patterns across the genome by preserving DNA methylation after replication events (Goll and Bestor 2005). DNMT methylates cytosines in a *de novo* fashion (Goll and Bestor 2005) and has been shown to respond to environmental stimuli (reviewed in Flores et al. 2013). Despite possessing all the conserved amino acid residues necessary for DNA methylation, DNMT2 proteins target and methylate tRNA molecules (reviewed in Jeltsch et al. 2017). Intriguingly, Dnmt2 is the only methyltransferase found in *Drosophila,* methylating DNA (Kunert et al. 2003) at uncharacteristically low rates for animals (Deshmukh et al. 2018). Dnmt2 is separated from the other DNMT clades phylogenetically (Jeltsch et al. 2017), though it does not group with classic RNA-methyltransferases. DNMT2 is the most widely distributed methyltransferase and is generally thought to be the first DNMT (Ponger et al. 2005). Evidence suggests that DNMT3 has undergone multiple duplication events in several taxa (Del Castillo Falconi et al. 2022), resulting in paralogs like DNMT3a, b, or L. Despite the known relationships between and functions of the DNMTs, most studies do not study them jointly.

*Daphnia* are freshwater microcrustaceans that are used as a model organism in the fields of evolution, ecology, genomics, phenotypic plasticity, and, more recently, epigenetics (Vandegehuchte and Janssen 2009; Weider and Pijanowska 1993; Colbourne et al. 2011; Stollewerk 2010; Schield et al. 2016; Hearn et al. 2018, Nguyen et al. 2021, Agrelius et al. 2022). *Daphnia* studies show high levels of phenotypic plasticity in response to a variety of biotic and abiotic environmental cues (Garbutt and Little 2014; Jeremias et al. 2018) both within-and across-generations (Agrawal et al. 1999, Walsh et al. 2014, 2015; Agrelius et al. 2022; Sun et al. 2023). Changes in gene expression, DNA methylation, and protein structure have been observed several generations removed from the original cue experienced by the mother (Schwarzenberger and von Elert 2013; Ortiz-Rodriguez et al. 2012; Hales et al. 2017; Schield et al. 2016; Hearn et al. 2018).

*Daphnia* are cyclical parthenogens, usually reproducing asexually (Zaffagnini 1987; Harris et al. 2012), making *Daphnia* an ideal model system for studying transgenerational phenomena that may be epigenetic in origin. Furthermore, the publication of high-quality genomes (Coulborne et al. 2011; Ye et al. 2017) and the advances in gene expression control techniques like viral transgenesis (Robinson et al. 2006), CRISPR (Nakanishi et al. 2014; Ismail et al. 2018), TALEN (Nakanishi et al. 2015), and RNA interference (Kato et al. 2011; Hiruta et al. 2013; Schumpert et al. 2015) increase the versatility and power of the *Daphnia* system.

The *Daphnia* genome contains all three DNMTs, with at least two DNMT3 paralogs (Nguyen et al. 2021), and *Daphnia* have been shown to alter DNA methylation within-and across generations (Vandegenhuchte et al. 2009a and b, Jeremias et al. 2018, Trijau et al. 2018, Hearn et al. 2019, Nyguen et al. 2020). Furthermore, DNMT expression can be influenced by dietary restriction and is subject to maternal influence in clonal lineages (Agrelius et al. 2022). Recent studies have shown that *Daphnia* respond to environmental stressors, altering the DNA methylation of their offspring in a stable, heritable manner relative to the current environment (Feiner et al., 2022). This transmission could alter the phenotypic trajectory of offspring as an anticipatory maternal effect, examples of which are common in *Daphnia* (Agrawal et al. 1999; Garbutt and Little 2014; Plaistow et al. 2015; Agrelius & Dudycha 2025).

RNA interference (RNAi) is a post-transcriptional gene silencing mechanism that operates through sequence-specific cleavage of endogenous messenger RNA (mRNA) transcripts. The RNAi pathway is highly conserved across eukaryotes and has been used for loss-of-function studies in various model systems (Ketting 2011). The RNAi pathway is activated by double-stranded RNA (Ipsaro and Joshua-Tor 2015), complementary to the mRNA of a gene of interest. The dsRNA is targeted and cleaved into 21-24nt, single-stranded, short-interfering RNAs that act as guides for the degradation of complementary mRNA transcripts but can also induce unintended changes in nontargeted proteins (Scacheri et al. 2004).

RNAi methods for gene knockdowns are well-established for several organisms (Mutti et al. 2006; Drake et al. 2012; Li-Byarlay et al. 2013), including *Daphnia* (Kato et al. 2011; Hiruta et al. 2013; Schumpert et al. 2015). Most RNAi mechanisms require microinjections as the vector delivery method. This presents a logistical problem for the *Daphnia* system for three reasons: 1) *Daphnia* mortality increases rapidly when not immersed in water, 2) microinjections likely need to occur in an early-stage embryos which are a fraction of the size of an adult, 3) early embryos (stage 1) have hypertonic pressure and burst when pierced. Experimentally successful microinjections result in a permanent reduction or loss-of-function of the gene of interest, limiting the scope of the study. Schumpert et al. (2015) developed a novel and efficient method to achieve RNAi through a daily bacterial feeding regime that mitigates the logistical issues and allows for the return of function of the gene of interest by stopping the RNAi treatment.

The utility of a technique that allows for the return of function is in its ability to permit age-specific gene knockdown of most genes, including those that might prove lethal if knocked down during early developmental stages. DNA and histone methylation are essential elements for controlling gene expression, regulating developmental cycles, and cell fate determination (Jaenisch and Jahner 1984, Ng and Bird 1999, reviewed in Burgess 2014). Loss of a key epigenetic regulator like DNA methyltransferase 1 (Dnmt1) can lead to increased cell (O’Neil et al. 2018) and embryonic mortality (Liao et al. 2015), making knockout studies increasingly difficult.

We previously found significant differences in Dnmt expression between two clones of *D. pulex,* both within-and across-generations, and significant maternal effects in response to food limitation (Agrelius et al. 2022). Data connecting differential DNA methylation to environmental stimuli are mounting (eg. Trijau et al. 2018; Jeremias et al. 2018; Hearn 2021), but few studies have investigated the loss-of-function of the epigenetic regulators themselves (Nguyen et al. 2021). We aim to use the RNAi feeding regime to individually knockdown the expression of three Dnmts in the *D. pulex* clones MORG 5 and TRO 3. By using the RNAi feeding regime, we hypothesize that DNMT expression can be reduced to negligible levels without incurring high mortality rates. This would allow for mechanistic investigations into transgenerational epigenetic changes.

## Methods

### Design and Construction of RNAi Vectors

Primers were designed to amplify 300bp products from each DNMT gene to be used in RNAi vectors. In an attempt to minimize off-target effects, potential amplicons were aligned to mRNA sequences from the other DNMT genes. Primer sets were then selected only if alignments were less than 12 consecutive nucleotides between the amplicon and non-target DNMT mRNAs (Supplemental Figures 1, 2, and 3), and amplicons did not overlap with regions used for qPCR. Amplicon size was chosen to maximize the number of siRNAs produced from a single dsRNA product (300bp double-stranded RNA cut into ∼25nt single-stranded siRNAs totaling 24 potential guides for the RNAi machinery), generating a more robust RNAi response. Restriction enzyme sites for XbaI and NheI were engineered into the 5′ ends of selected primers to allow for subcloning between pGEM-T Easy and L4440 plasmids (Fire et al. 1998).

Experiment 1 targeted two locations on DNMT1 (Figure 1), while all work in Experiment 2 targeted one unique location in each of the three DNMTs (Figure 2). Selected amplicons were cloned into pGEM-T Easy plasmids (Promega) and verified by sequencing. pGEM-T Easy plasmids are designed with 3′-T overhangs that allow unmodified PCR products with poly-A tails to be cloned directly into the plasmid. Transformations were carried out using competent DH5-α *E. coli* cells (New England Biolabs), and plasmid purifications were performed using a Qiagen Plasmid Miniprep kit. DNMT amplicons were cut out of pGEM-T Easy constructs using appropriate restriction enzymes and gel-purified for subcloning into L4440. Finally, glycerol stocks were made using 500μL of transformed DH5-alpha bacteria with positive recombinant pGEMT Easy constructs and 500μL of glycerol. Stocks were stored at -80°C.

**Figure 1.**
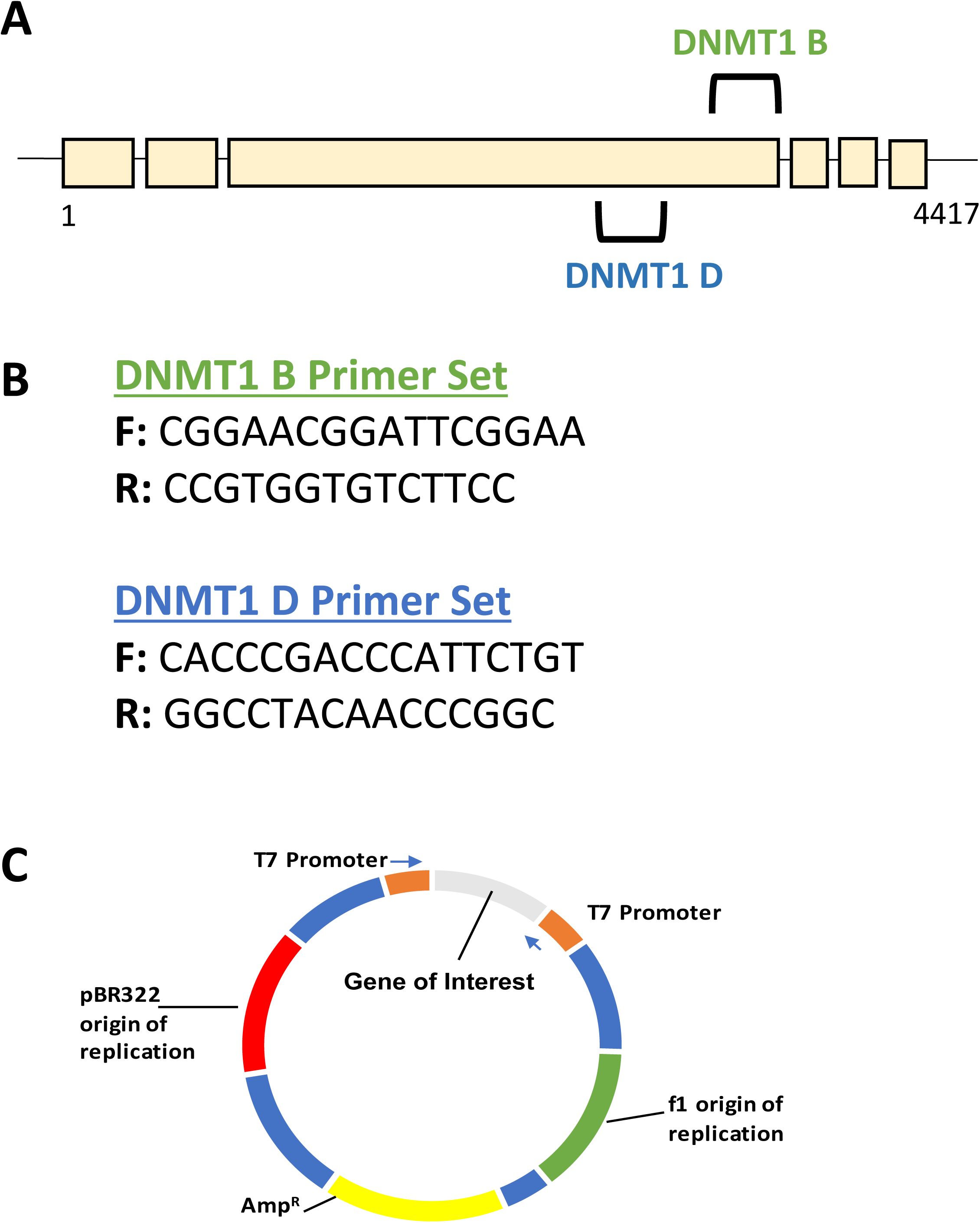
**A)** *Daphnia pulex* DNMT1 gene schematic. Marked on the schematic are the two regions amplified by PCR and cloned into pGEMT-Easy and L4440. **B)** Primer sets used to amplify 300 nucleotide regions of the DNMT1 gene. There is no overlap between the two amplicons. **C)** Schematic diagram of the L4440 plasmid vector designed by Fire et al. 1998. The flanking T7 promoters (orange) can be induced to produce double-stranded RNA of the cloned insert from the gene of interest (gray).

**Figure 2.**
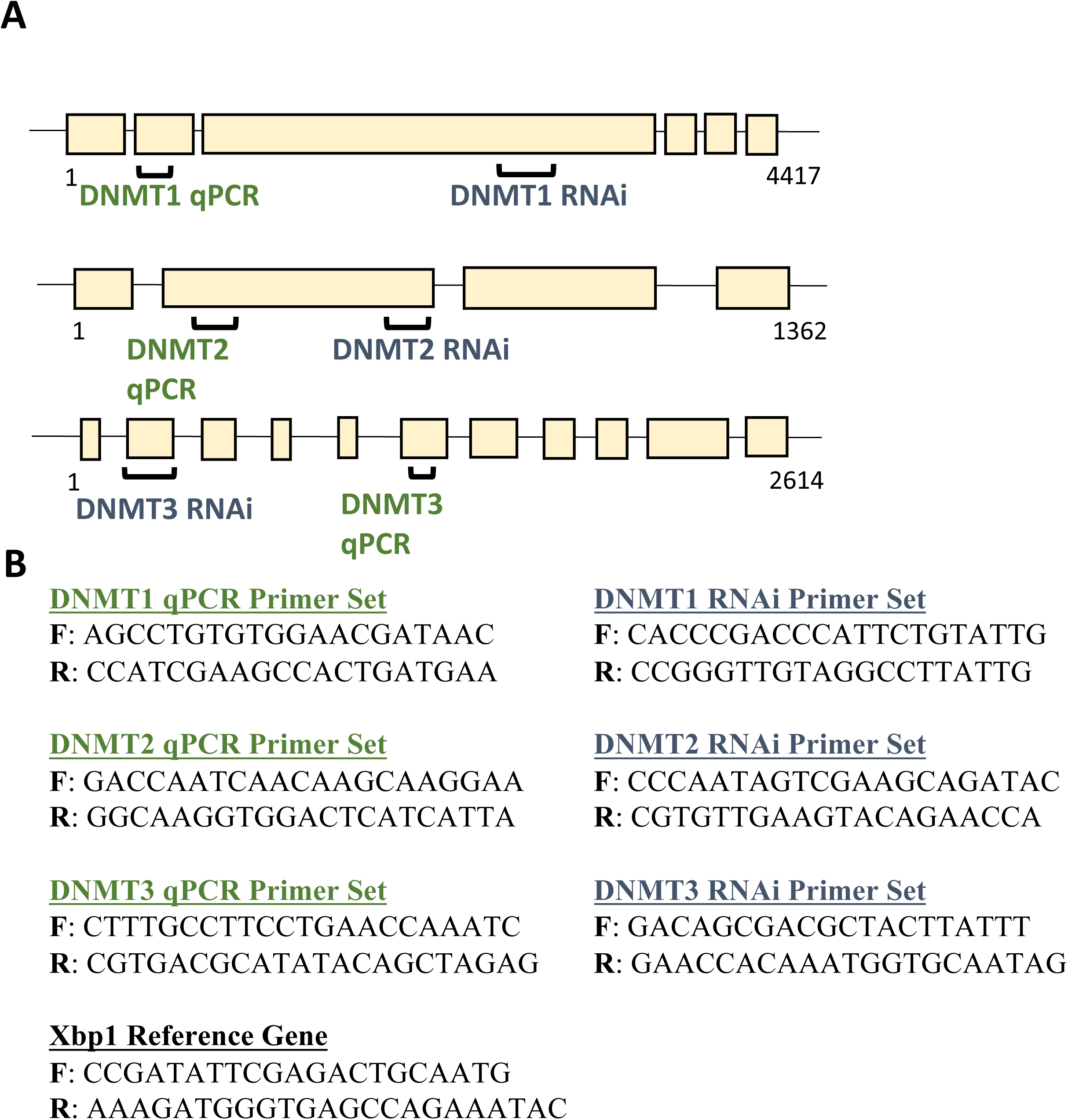
**A)** *Daphnia pulex* DNMT1, 2, and 3 gene schematics. Marked on the schematic are the two regions amplified by PCR for cloning into L4440 (RNAi) or qPCR. **B)** Sequence information of the primer sets used to amplify 300 nucleotide regions of the DNMT genes (RNAi) and 100 nucleotide regions (qPCR). There is no overlap between the two amplicons. Xbp1 was used as a reference for gene expression analyses.

Purified DNMT PCR products were subcloned into L4440 plasmid vectors (L4440 Plasmid: 1654, gifts from Dr. A. Fire to Addgene). The L4440 vector allows for cloning PCR products between two T7 promoters oriented in opposing positions (Figure 1). All L4440 plasmids were treated with Calf Intestinal Phosphatase (CIP, New England Biolabs) prior to ligation reactions to increase ligation efficiency. Cell transformations were performed using HT115 (DE3) (W3110, rnc14::DTn10 (Addgene; Dasgupta et al. 1998) *E. coli*. HT115 bacterial strains contain a source for T7 RNA polymerase (DE3 lysogen), which can be induced through the addition of 2mM IPTG and are deficient in RNase III, leading to efficient production of dsRNA of the amplicon cloned between the two T7 promoters of L4440.

We set up a bacterial feeding regime (Schumpert et al. 2015) that exploits the RNAi pathway to induce endogenous gene knockdown in the *Daphnia* system. Transformed bacteria (bacteria with the plasmid L4440 containing an amplicon from a gene of interest) were grown overnight (∼14-16hours) in Luria Broth (LB) with 2mM of IPTG to induce the production of dsRNA. The optical density (OD_600_) of bacterial cultures was measured using an Eppendorf BioPhotometer. Bacterial cultures from an OD_600_ of 2.8 units were pelleted and resuspended in 200mL of filtered lake water to feed the bacterial culture to *Daphnia*, resulting in a final concentration of 0.014 OD_600_ or ∼1.2*10^7^ *E. coli* cells. New bacterial cultures were prepared daily using fresh LB and IPTG. In addition to the bacterial cells, *Daphnia* were fed quantitatively 20,000 cells/ml of *A. falcatus* daily, resulting in a mixture of bacteria and algae. New bacterial cultures were prepared daily throughout their administration to *Daphnia*.

### Testing Effects of DNMT RNAi Vectors

Two experiments were conducted using the RNAi vectors and bacterial feeding regime. Experiment 1 used two *D. pulex* clones, MORG 5 and TRO 3, and two RNAi vectors that targeted DNMT1. Experiment 2 targeted all three DNMT genes with unique RNAi vectors using the MORG 5 clone. The *D. pulex* clones originated from temporary ponds in Maine and Michigan, USA, respectively, and have been cultured under constant laboratory conditions for many years. All animals were propagated using filtered lakewater (to 1µm) collected locally from an outflow of Lake Murray in Lexington County, SC, USA, and maintained at 20°C under a 12:12 light:dark photoperiod. Laboratory-reared animals were fed a daily diet of vitamin-fortified *Ankistrodesmus falcatus* (Goulden and Horning 1980). To minimize potential maternal effects-related variation in gene expression and epigenetic motifs (Agrelius and Dudycha, 2025), three acclimation generations were reared under standardized, high food conditions. Animals in each generation were taken from the third clutch and fed a daily diet of 20,000 cells/mL of *Ankistrodesmus falcatus.* Each animal was housed separately in a 150mL beaker with 100mL of filtered lake water and transferred every other day into a new beaker with fresh lakewater. Each beaker was lightly dusted with cetyl alcohol to prevent entrapment on the water surface (Desmarais 1997).

Experiment 1 consisted of replicate cohorts of 10 female *Daphnia* per clone in 250mL beakers with 200mL of filtered lakewater. Eight replicate cohorts were randomly assigned to one of three treatments: DNMT1B, DNMT1D, or Empty Vector (EV). DNMT1B and DNMT1D are RNAi vectors targeting different regions of DNMT1 mRNA transcripts (Figure 1). For the MORG 5 DNMT1B treatment, we could only set up 5 replicate cohorts due to the lack of female neonates at the time of setup. Offspring born to females in each vector treatment during the experiment were collected and set up in cohorts of 10 under the same feeding regime as their mothers.

Experiment 2 consisted of six treatments: three controls, and three RNAi DNMT treatments, which can be seen in Table 1. Three controls were necessary to 1) serve as a negative control and 2) partition out the effects of bacterial and dsRNA presence on DNMT gene expression. Replicate cohorts of 10 *Daphnia* per treatment in 250mL beakers with 200mL of filtered lakewater were randomly assigned to one of the six feeding treatments (total of n=870 experimental animals). Most treatments consisted of 15 replicates but we could only establish 12 replicate cohorts for the Algae-only control due to the lack of births during the experimental setup.

**Table 1:** List of Experiment 2 Treatments

| **Treatment** | **Information on and purpose of the treatment in the experiment** |
| --- | --- |
| *Algae*<br>**Control** | Negative control, *ad libitum*, or high food diet consisting of 20,000 cells per mL of *Ankistrodesmus falcatus* algae. |
| *Empty Vector (EV)*<br>**Control** | Isolate the effects generated by increased bacterial concentrations on *Daphnia* life history and DNMT expression. Empty L4440 vector used to transform Ht115 *E. coli* cells. |
| *GFP-dsRNA (GFP)*<br>**Control** | L4447 plasmid with 5' half of the GFP ORF used to isolate the effects generated by the presence of double-stranded RNA species on life history and DNMT expression. GFP is a nonendogenous target, any effect observed would be attributed to the RNA species and or increased bacterial concentrations. |
| *D1RNAi (D1)* | L4440 plasmid with the PCR product from DNMT1; knockdown the expression of DNMT1 |
| *D2RNAi(D2)* | L4440 plasmid with the PCR product from DNMT 2; knockdown the expression of DNMT2 |
| *D3RNAi(D3)* | L4440 plasmid with the PCR product from DNMT 3; knockdown the expression of DNMT3 |

Bacterial feeding treatments for both experiments were initiated within 5-8 hours of animals’ maturation (defined as eggs being deposited into the brood chamber) and continued for 10 days. We recorded mortality, the number of offspring produced per day (or time to maturation for the offspring in Experiment 1), and several qualitative observations pertaining to animal health during the 10-day bacterial feeding regime. Due to complications incurred from COVID-related shutdowns, animal tissue for qPCR gene expression work could not be collected for Experiment 1. Animal tissue was collected at the end of the 10-day trial in Experiment 2 and stored in RNAlater (Invitrogen) at 4C for four days before being stored at -80C until RNA extractions.

### Gene Expression

For total RNA extractions, animals were pooled into cohorts (10 animals per treatment) for a single biological replicate. Prior to extraction, eggs were carefully dissected out of the brood chamber to exclude embryonic gene expression. Total RNA was then extracted from homogenized tissue using a trizol/purezol-chloroform method (Purezol, Bio-Rad). Sample purity, concentration, and integrity were assessed using Nanodrop 2000, Quibit, and gel electrophoresis. gDNAase Iscript (Bio-Rad) was used to remove any genomic contamination and convert into first-strand cDNA.

Gene-specific qPCR primers were previously designed and validated for all three DNMTs (Agrelius et al. 2022). Primer sequences and relative positioning compared to amplicons used for RNAi can be seen in Figure 2. qPCR primers amplified nonoverlapping regions of each gene to ensure dsRNA produced by the plasmid vector treatment would not be detected. qPCRs were run with three technical replicates per biological sample using PowerUp SYBR Green Master Mix (Applied Biosystems) in a BioRad CFX96 Real-Time PCR System (BioRad).

Thermal cycling conditions were: 2 min at 50°C and 2 min at 95°C, followed by 40 cycles of 15 sec at 95°C and 30 sec at 60°C. Dissociation curve analysis and gel electrophoresis were performed to confirm correct amplicon size and primer specificity. No-template controls (NTC) and no-reverse-transcriptase (NRT) controls were used to confirm the absence of contamination. Relative DNMT mRNA levels were normalized to XBP1 transcript level using the Pfaffl method, with an efficiency correction calculated from Real-time PCR Miner (Zhao et al. 2005).

## Statistical Analyses

Statistical analyses were conducted in R, version 3.6.2 (R Core Team 2021), and plots were generated using ggplot2 (Wickham 2016). Effects of the RNAi treatments on survivorship (*l_x_*), a type of ‘time to event’ data, was analyzed using a Cox proportional hazards regression with an efron method to handle tied events. To test for the effects of each RNAi treatment on offspring production, the total number of offspring produced (per treatment) was divided by the total number of surviving mothers (per treatment) for each day. RNAi treatment effects on offspring data were analyzed using a type-III ANOVA. Effects of RNAi bacterial treatments on gene expression data were analyzed using a Kruskal-Wallis rank-sum test and a Dunn’s Test for post hoc multi-comparisons. Data normality was tested using Shapiro-Wilks tests and visual comparisons; the number of offspring was log-transformed to meet the assumptions of normality. Data variance heterogeneity was assessed using Levene’s Test.

## Results

### Experiment 1

MORG 5 survivorship was not significantly reduced by Dnmt1B or Dnmt1D RNAi vector treatments compared to animals fed bacteria with empty vectors, Figure 3 (χ^2^ = 0.142, p = 0.931, *df* = 2). TRO 3 survivorship was significantly reduced by RNAi vector treatments (χ^2^ = 60.974, p < 0.0001, *df* = 2), with the Dnmt1B RNAi vector increasing the probability of death by three times that of the empty vector (Fig. 3; p *<* 0.0001, *hazard ratio* = 3.765, *CI* = 2.694 -5.260). The number of offspring produced per living female was reduced in all three treatments (DNMT1 B/D; EV) in both clones when compared to the maternal generation (MORG 5: F_3,23_ = 4.506, p < 0.012; TRO 3: F_3,13_ = 9.479, p < 0.001) but not between RNAi vector treatments (Supplemental Fig. 4). Fecundity did not significantly differ between MORG and TRO in the maternal generation, G_0_, used to produce experimental animals, G_1_.

**Figure 3.**
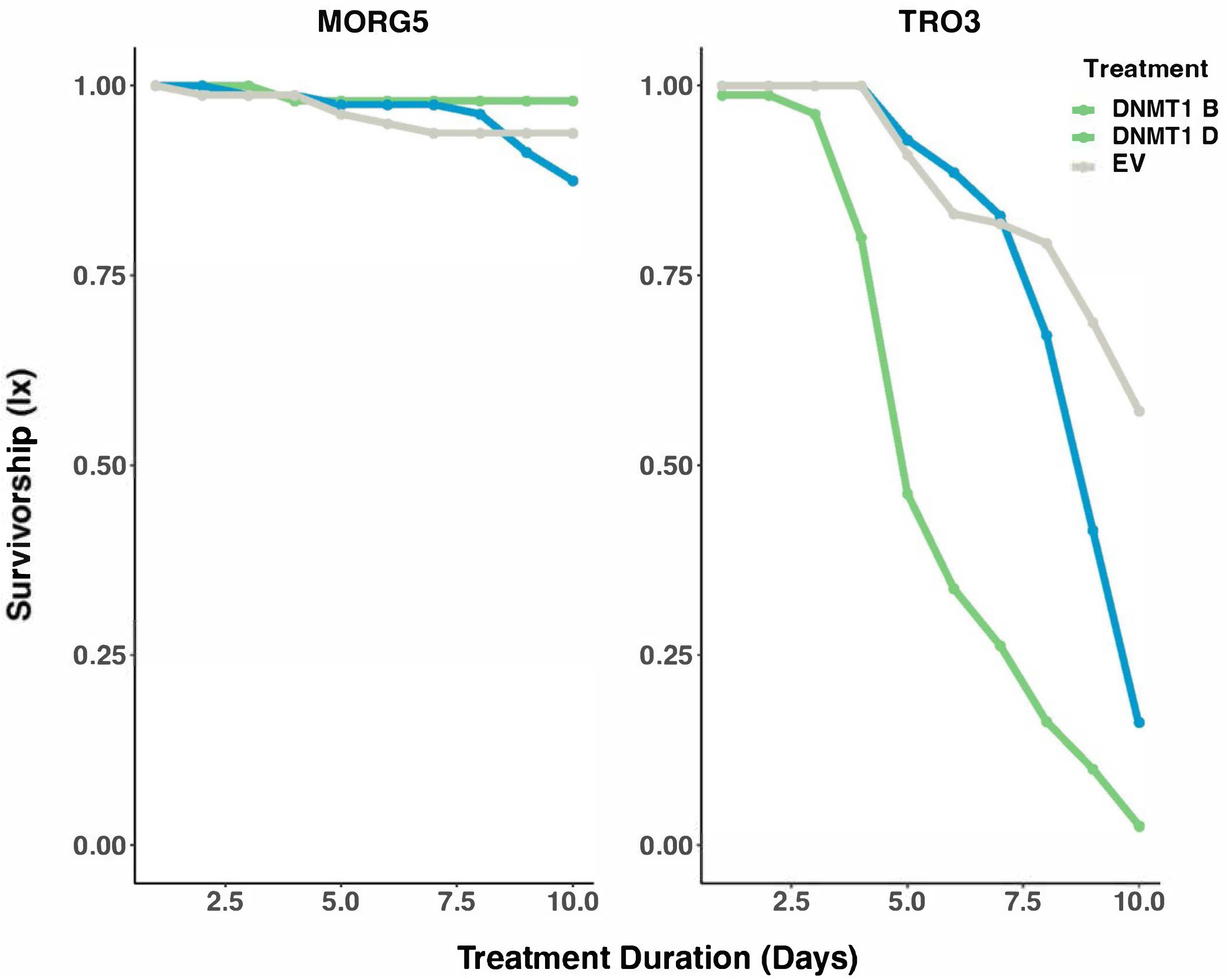
Plots showing the effects of each vector type on survivorship (lx) of the two clones, MORG5 and TRO3. Animals treated with the DNMT1 B vector are shown in green, DNMT1 D vector in blue, and EV in gray. No significant difference was observed between either DNMT vector when compared to EV in MORG5 (𝜒^2^ = 0.142, p = 0.931, *df* = 2). 98% of MORG5 DNMT1 B, 76% of DNMT1 D, and 94% EV survived the 10-day trial. DNMT1 treatment significantly reduced survivorship in TRO3 when compared to EV (𝜒^2^ = 60.974, p < 0.0001, *df* = 2). Death was 3 times as likely to occur when treated with DNMT1 B (p < 0.0001, *HR* = 3.765). There was no significant increase in mortality between DNMT1 D and EV (p < 0.0966, *HR* = 1.328). 2.5% of TRO3 DNMT1 B, 16% of DNMT1 D, and 60% of EV survived the 10-day trial.

Photographs were taken after ten consecutive days of the RNAi vector treatment (Supplemental Fig. 5). Notable differences in animal body and gut coloring, presence or absence of vitellogenesis, offspring production, and carapace abnormalities were observed between the two genotypes. Note, we had no anticipated observing these kinds of characteristics and thus our observational comparisons are informal because the photographs were not standardized. For all three RNAi vectors, gut coloring ranged from dark brown to clear in TRO 3 animals. Surviving animals from the Dnmt1D RNAi vectors developed a “curly” phenotype in which deformities in the carapace impede or curl into the brood chamber. Vitellogenesis was not observed for the majority of the TRO 3 animal cohorts. In contrast, MORG 5 animals generally appeared less translucent, with gut coloring ranging from a vibrant green to a bright yellow. Vitellogenesis was commonly observed among all three vector treatments, and embryo development was generally unimpeded by the bacterial feeding regime.

Neonates, G_2_, produced during Experiment 1 vector treatments from both clones were collected and treated with the same RNAi vector as their mother, G_1_. In both DNMT RNAi vector treatments, neonates experienced at least 50% mortality before reaching maturation. All TRO 3 offspring that survived developed a brood chamber (Supplemental Fig. 6) but did not begin the process of vitellogenesis during a 12-day bacterial feeding regime. Body pigmentation was within a normal range, and gut coloring remained a vibrant or pale yellow-green (Supplemental Fig. 6). MORG 5 neonates from the Dnmt1D and EV treatments were observed to have completed the process of vitellogenesis on the 10^th^ and 11^th^ day of the 12-day trial. Embryo development appeared normal. Pigmentation and gut coloring resembled that of normal, well-fed *Daphnia*. Fat deposits were observed in neonates from both clones.

Due to the adverse effects induced by the vectors on the TRO 3 clone in Experiment 1, we only used the MORG 5 clone to examine the potential effects of DNMT RNAi further. We did, however, test the effects of nonendogenous dsRNA on TRO 3 on survivorship in Experiment 2 (Supplemental Fig. 7). GFP-dsRNA (GFP) vectors significantly reduced TRO 3 survivorship (χ^2^ = 14.046, p < 0.029, df = 6), with only ten of thirty TRO 3 animals surviving a 10-day trial. Remaining TRO 3 animals appeared translucent, with no pigmentation in the carapace outside the compound eye and little to no coloring in the gut (Supplemental Fig. 8B). Despite reaching maturation before the experiment’s start, no offspring were released from the brood chamber. Additional vitellogenesis events were not observed during the GFP bacterial treatment.

### Experiment 2: RNAi Vector Effects on Phenotypic Traits

MORG 5 survivorship was not significantly reduced by RNAi vector treatments (χ^2^ = 0.366, p < 0.996, df = 5; Supplemental Fig. 9). No significant difference in the number of offspring per female was detected between the MORG 5 RNAi vector treatments (F_5, 42_ = 0.024, p = 0.997, df = 5; Fig 4).

**Figure 4.**
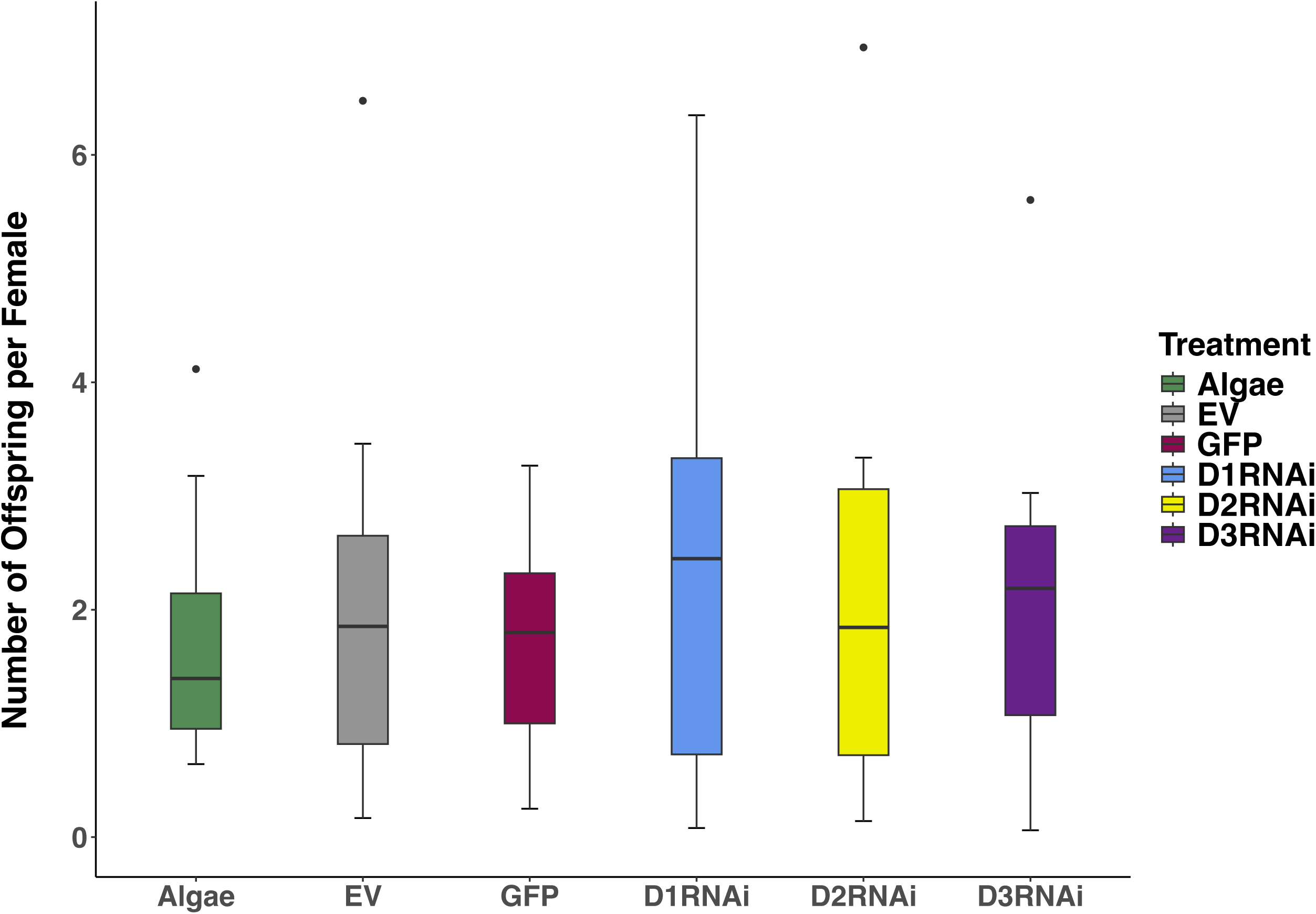
Boxplots of the number of offspring produced by the MORG5 clone. Values were standardized by dividing the total number of neonates produced each day by the total number of surviving females. Treatment identification is as follows: Algae (green), Empty Vector (grey), L4440 with GFP (turquoise), and each DNA methyltransferase RNAi treatment 1 (blue), 2 (yellow), and 3 (purple). No significant difference was detected between treatment (F_5, 42_ = 0.024, p = 0.997, *df* = 5). TRO3 animals in the GFP plasmid vector did not produce offspring during the 10-day bacterial feeding regime and is not included in the figure.

Within 18 hours of receiving the first vector dose, MORG 5 animals in the D1RNAi, D2RNAi, and D3RNAi treatments prematurely shed their carapace. After the second vector dose, animals from the same treatments molted again. We did not observe similar events in the Algae, EV, or GFP controls. Photographs from each treatment on day 10 showed distinct differences in animal pigmentation, gut coloring, and reproduction (Supplemental Fig. 8).

MORG 5 animals from the Algae, EV, and GFP treatments showed normal pigmentation and gut coloring ranging from pale to bright green, indicating continuous feeding on *A. falcatus* (green algae) and presumably bacteria. Animals from Dnmt RNAi treatments showed abnormal pink pigmentation, suggesting elevated levels of hemoglobin. Gut coloring appeared a bright green. Continued vitellogenesis was observed for all MORG 5 animals, and embryo development appeared normal.

### Experiment 2: RNAi Vector Effects on Gene Expression

DNMT expression in the three controls used for Experiment 2 (Table 1) was scaled to the Algae control cohort (Fig. 5). Significant increases in DNMT expression were observed in both EV (DNMT2 and 3) and GFP (DNMT 1, 2, and 3) treatments (Tables 2, 3, 4). EV treatments increased DNMT2 and 3 expression by ∼0.3 fold; GFP treatments increased DNMT1, 2, and 3 expression by 1.47, 0.7, and 1.59 fold, respectively. DNMT expression was significantly reduced by DNMT RNAi vector treatments when scaled to the GFP control (Fig. 6; Supplemental Tables 1, 2, and 3). However, DNMT RNAi vector treatments did not reduce DNMT expression levels below the Algae control. We observed off-target DNMT expression reductions in each DNMT RNAi vector treatment (Fig. 6). The magnitude of the off-target reduction varied between RNAi treatments (Fig. 7).

**Figure 5.**
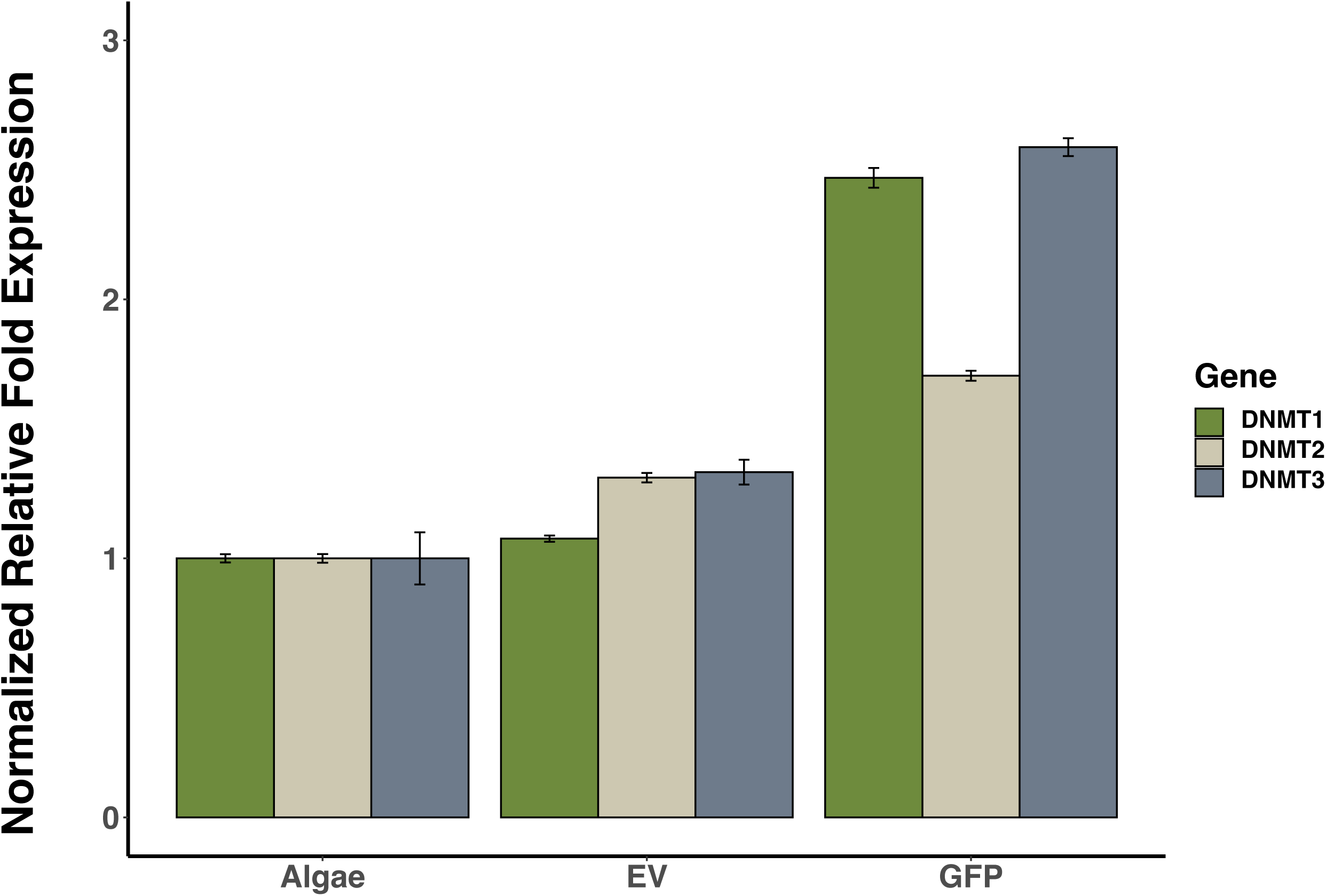
DNMT expression relative to XBP1 and normalized to the Algae control. EV treatments significantly increased DNMT2 and 3 expression. dsRNA (GFP) significantly increased mRNA transcripts for all three methyltransferase genes.

**Figure 6.**
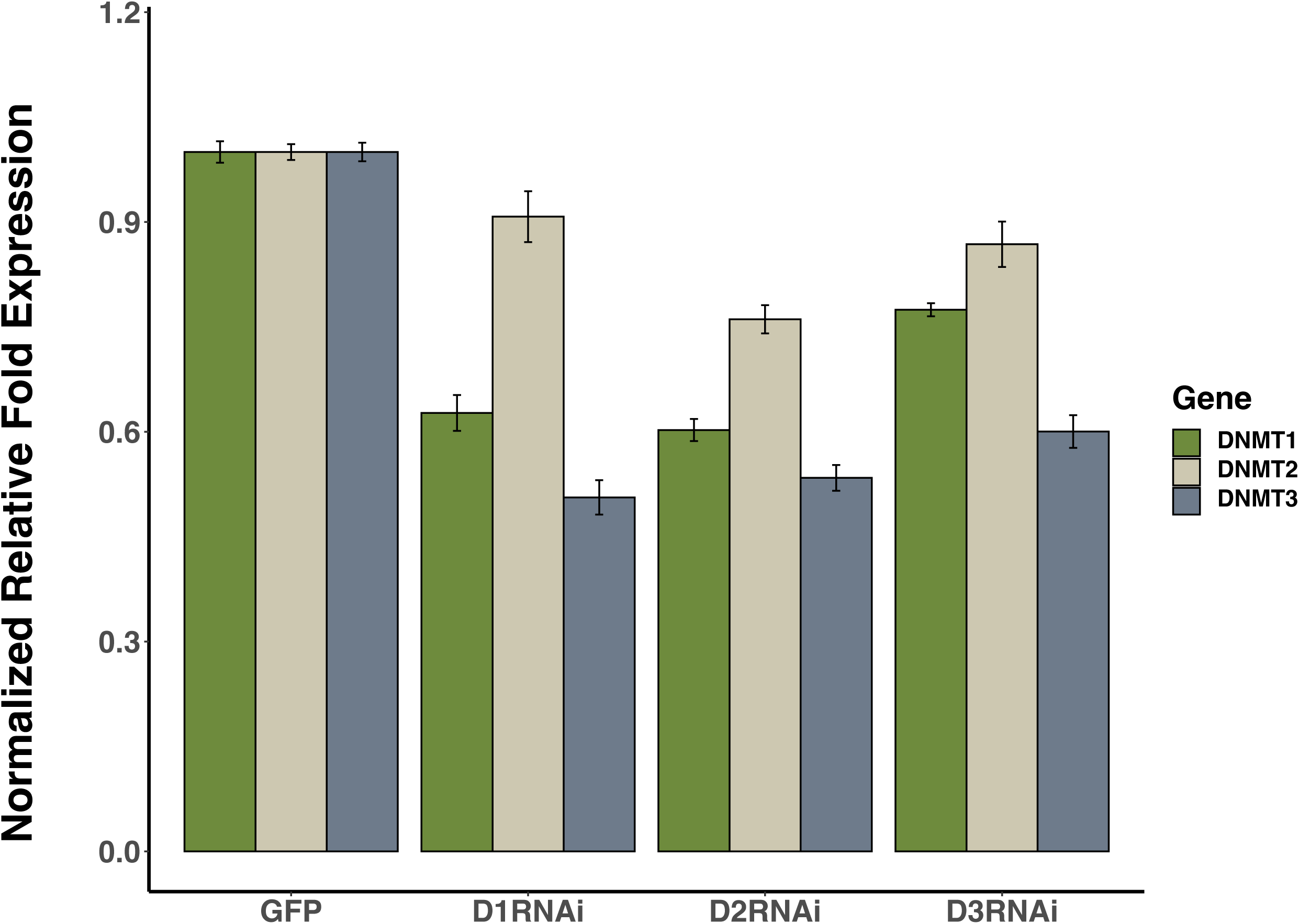
DNMT expression relative to XBP1 and normalized to the GFP control. Note that names for each DNMT RNAi vector treatment were abbreviated to conserve space. Nomenclature is as follows: D1 refers to the DNMT1 RNAi vector treatment. Expression of nontarget methyltransferase genes also reduced in each RNAi treatment.

**Figure 7.**
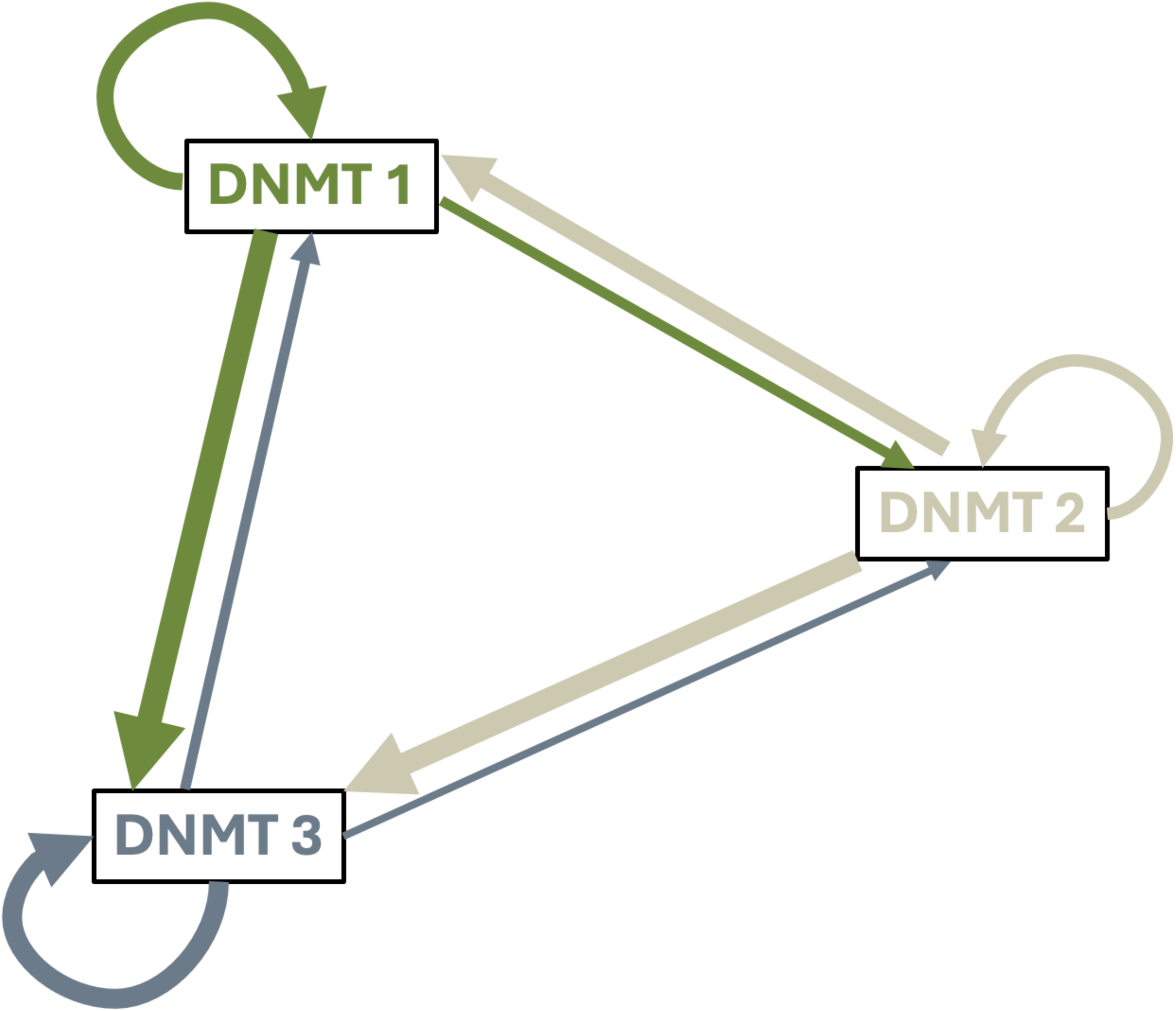
Magnitude of on-and off-target expression reduction indicated by line thickness. Curved arrows indicate that DNMT RNAi vector treatment on the DNMT of interest. Straight arrows indicate the direction of the off-target effect of each DNMT RNAi vector treatment on other DNMTs.

## Discussion

We found significant impacts of the L4440/HT115 RNAi bacterial feeding regime on DNA methyltransferase (DNMT) gene expression and *Daphnia* biology. Although we were unable to measure gene expression in our first experiment due to COVID-related restrictions, it helped us focus our second experiment on a single *D. pulex* clone that tolerated the RNAi procedure well. That experiment produced three intriguing findings regarding RNAi treatments and DNMT gene expression. First, DNMT expression in *D. pulex* clone MORG 5 significantly increased in response to being fed either the empty vector or the vector with the GFP RNAi construct compared to being fed only the algae control. Second, all three DNMT RNAi treatments significantly reduced DNMT expression compared to GFP RNAi. Third, targeting any one DNMT using RNAi resulted in a reduction of mRNA transcripts of the other two DNMTs, although the off-target effect differed in magnitude.

Observing expression changes in DNMT3 and DNMT1 following exposure to the empty vector and GFP RNAi treatments was not unexpected. The greater than two-fold expression increase in the GFP RNAi treatment suggest these genes are substantial parts of the *Daphnia* response to the presence of double-stranded RNA. DNMT3 is a *de novo* methyltransferase capable of responding to environmental cues (Goll and Bestor 2005), while DNMT1 functions as a maintenance methyltransferase, preserving DNA methylation during replication events (Goll and Bestor 2005). Pathogen exposure and infection, including *E. coli,* have been shown to induce and repress DNA methyltransferase activity (Qin et al. 2021) and induce differential methylation patterns across thousands of loci, resulting in extensive epigenetic remodeling and changes in gene expression (Pacis et al. 2015; Zhou et al. 2024) capable of influencing immune responses (Tolg et al. 2011).

Double-stranded RNA immune responses are very well characterized in vertebrates to induce gene silencing using an antiviral program mediated by Type 1 interferons (Robalino et al. 2004) that have pleiotropic functions in both innate and adaptive immunity. Invertebrate genomes lack genes associated with the interferons; however, Robalino et al. (2005) observed a robust antiviral immunity induced by dsRNA in marine shrimp and sequence-specific targeting and destruction of mRNA transcripts. Unintended consequences like stimulation of immune system responses (Robalino et al. 2005), and nonspecific protein targeting via the siRNA produced in the RNAi pathway are well documented in other systems (Fellman and Lowe 2014; Kaelin 2012, reviewed in Meng and Lu 2017, Knudesen-Palmer et al. 2024).

RNAi relies on triggering an innate immune response that destroys complementary nucleic acid targets. Our targets were mRNA transcripts from genes responsible for maintaining the stability of the genome through DNA methylation (Bonasio et al. 2010). However, the elevated concentration of transformed bacteria compared to the algae-only control may have initiated a separate immune response. Bacterial-mediated immune responses in *Daphnia* rely on cell surface receptors from the TOLL gene family that function as signal transducers (McTaggart et al. 2009; TOLL reviewed in Duan et al. 2022). TOLL proteins show specificity with interactions between bacterial strains and rely on several well-conserved genes for signal transduction (McTaggert et al. 2009, references within). Orthologues of at least seven of those genes have been identified in *D. pulex* (TOLL, Myd88, Relish, Pelle, Cactus, Imd, STAT) and other immune-related genes that respond directly to bacteria (Michel et al. 2001). The TOLL pathway has been identified as a vital component in the *Drosophila* antiviral response (Zambon et al. 2005; Wang et al. 2014; Chauhan et al. 2025), and can be stimulated by *E. coli* as well as dsRNA in crustaceans (Robalino et al. 2004, references within). Immune responses to pathogens in *Daphnia* have been detailed as early as 1884 (Metchnikoff 1884), making it plausible that the responses observed in our studies are due to an immune response outside the traditional RNAi-directed pathway.

### Off-Target Effects of RNAi Knockdown of Dnmts: Nonspecific DNMT Knockdown

Though extensive measures were taken to ensure that our RNAi constructs would target transcripts from only the gene of interest, we observed a reduction of non-targeted DNA methyltransferase expression in all three of our DNMT RNAi treatments. The magnitude of the off-target expression reduction varied based on which DNMT was targeted (Figure 7) and which off-target DNMT was observed. Crosstalk between epigenetic machinery is heavily documented (Kan et al. 2022; Lempiainen et al. 2023), resulting in changes in the expression of multiple epigenetic genes. The targeted reduction of one DNMT could impact the expression of another as part of DNMT biology, or an alternative mechanism could be facilitating the off-target reduction.

Within the RNAi machinery, the argonaute protein is guided by 20-24nt siRNAs to cleave mRNA targets (reviewed by Faehnle and Joshua-Tor 2007). We ensured that each 300nt methyltransferase amplicon cloned into L4440 vectors had no more than a 12nt continuous alignment with mRNA from the other methyltransferase genes. This should eliminate the possibility of cross-reactivity mediated by the argonaute protein in the RNAi complex due to potential siRNAs lacking specificity. However, it is possible that the imperfect matches of the siRNA resulted in off-target translational repression and subsequent mRNA decay that is argonaute-independent. Endogenous microRNAs (miRNAs) can lead to translational repression by binding to mRNA, thereby preventing translation and facilitating exonucleolytic decay (reviewed by Neumeier and Meister 2021). Only partial complementarity is needed between the miRNA and mRNA for the translational repression to occur (Saxena et al. 2003). This mechanism could explain the nonspecific reduction of methyltransferase gene expression.

### Insight into Daphnia Biology: RNAi effects on Fecundity, Phenotype, and Genotype

We expected to observe some genotypic differences between the two clones during Experiment 1. Previous work with the two clones had shown TRO3 to delay maturation, switch to resting egg production, and increase DNMT expression in response to reductions in resource quantity (Agrelius et al. 2022). Still, the significant contrast in responses between the two clones was unexpected, especially in the empty vector treatment. Schumpert et al. (2015) did not observe changes in reproduction for either *D. melanica* or the *D. pulex* genotype used, nor were there mortality rates exceeding 20%. Subsequent *Daphnia* RNAi studies report effective gene knockdowns without unexplained mortality in *D. pulicaria* (Schumpert, et al. 2016), *D. magna* (Street et al. 2019), *D. pulex* (Lin et al. 2019; Zhou et al. 2020), and *D. galeata* (Cao et al. 2023). In fact, Lin et al. (2019) reported offspring production was increased in RNAi-treated animals compared to controls. Thus, it seems clear that the extreme susceptibility of the TRO3 clone to RNAi-by-feeding is not the norm in *Daphnia*.

Visible signs of stress were observed during Experiment 1 in the TRO3 animals. Signs included higher mortality rates, ranging from 40% to 97.5%, including the empty vector, where nearly half of the cohort died. We informally observed that all TRO3 animals reduced their filtering rates or stopped them completely, and their gut coloring shifted from green to brown. Filter-feeding in *Daphnia* is physically linked to respiration, with most of their oxygen supply coming from the internal current generated by the feeding combs’ movement (Pirow et al. 1999). By slowing or halting the filtering process, algal cells remain in the gut and decompose, resulting in a yellow or brown color; effectively, the animal loses the nutrient content of the food and starves while also experiencing hypoxia. This was observed in both DNMT1 and empty vector treatments of the TRO3 clone but not in MORG5. Rather, MORG5 seemed relatively unaffected by the vector treatments and continued producing offspring throughout the ten-day trial.

MORG5 animals in Experiment 2 experienced low mortality and showed no significant differences in offspring production among RNAi or control treatments (Figure 4), unlike the TRO3 animals, which produced no offspring when treated with GFP RNAi vectors. However, MORG 5 animals showed clear stress and potentially immune-related responses in all of the RNAi treatments (GFP and DNMT) that were not observed in the TRO3-GFP treatment, namely a startling double ecdysis event within 36 hours of the bacterial treatment and high levels of apparent hemoglobin production. The different responses of TRO3 and MORG5 show that *Daphnia pulex* harbors major genetic differences in how animals respond to double-stranded RNA, despite otherwise normal life histories.

Targeting methyltransferase transcripts could be a primary cause for the observed premature molting in MORG5. Molting in juvenile *Daphnia* is controlled primarily through the hormone 20-hydroxyecdysone and its interaction with juvenile hormones (Gilbert et al. 2002; Baldwin et al. 2001). Molting results in a major loss of calcium: ∼90% of total body calcium is lost each molt (Alstad et al. 1999). This loss is not replaced through diet, but rather, calcium uptake is an active process (Cowgill et al. 1986). Juvenile hormones respond to environmental cues, resulting in increased expression of hemoglobin (Gorr et al. 2006), changes in morphology (Miyakawa et al. 2013), and oogenesis (Charniaux-Cotton 1985). We began feeding bacteria six days after birth, which is the average date of maturation for MORG5 (Agrelius et al. 2022). At this early stage, developmental processes are still mediated through juvenile hormones and presumably susceptible to environmental cues. DNMT3 responds to environmental cues and establishes new methylation patterns across the genome, while DNMT1 maintains the existing methylation profile, especially during development (Goll and Bestor 2005). Both genes are highly expressed during developmental processes, and methylation profiles across the genome show differential patterns (Goll and Bestor 2005). Changes in expression for either methyltransferase during the first reproductive event could impact the ecdysis process, resulting in both the abortive event and molting being observed.

Our work provides an interesting and novel insight into DNMT and *Daphnia* biology. DNMT 1, 2, and 3 expression is rarely studied together, and our results suggest a potent crosstalk mechanism between the methyltransferases that warrants further investigation in joint studies. Future research can investigate several questions, including whether the RNAi effect on DNMTs can be fine-tuned to limit off-target effects. What is the relationship between RNAi, genotype, and mortality? Do the DNMT RNAi treatments alter DNA methylation or other epigenetic markers? Does RNAi influence maternal effects? Could the loss of methyltransferase activity limit plasticity, thereby reducing the variance observed in trait means responding to environmental stressors?

## Acknowledgments

Matt Bruner, Rachel Schomaker, and Jake Swanson assisted with various tasks during the work. The Morg-5 and Tro-3 clones were kindly provided to us by Mike Lynch. Funding for this work was provided by a grant from an NSF award DEB-1556645 to JLD and a University of South Carolina SPARC Graduate Research Grant awarded to TCA.

## Author Contribution Statement

TCA and JLD conceived the ideas and designed the experiments and led the writing of the manuscript; SB and TCA performed all cloning and recombinant work; TCA, AR, and KH collected the data; TCA analyzed the data; All authors contributed critically to the drafts and gave final approval for publication.

## Data Archiving

Data will be available in Dryad upon acceptance.

