## Supplemental File for "Genetic background influences the effects of RNAi-by-feeding for DNA methyltransferases in *Daphnia pulex*"

```
Dnmt1mRNA           GAAAAGTACAGGAAAACGGACAACATCAAGGGATCGAATAACGACACACCCGACCCATTC
Dnmt1amplicon       ----------------------------------------------CACCCGACCCATTC
Dnmt3amplicon       --------------------------------------------GACAGC-GACGCTACT
Dnmt2amplicon       --------------------------------------------CCCAATAGTCGAAGCA
                                                                  **   * *

Dnmt1mRNA           TGTATTGGCTATGTCGTTGGTATTGCCTATAATGGCA-TCCTGCACAACAATTTGA----
Dnmt1amplicon       TGTATTGGCTATGTCGTTGGTATTGCCTATAATGGCA-TCCTGCACAACAATTTGA----
Dnmt3amplicon       TATTTTTGTGGTTCGGATGACGATGATGGTGATGGAAATTTCGGAGTAGAAGCTGA----
Dnmt2amplicon       GATACTACCT-TGTAGCTAAAAAGTGTACAGACTTCTCATTTGGATCTGAAAACGACATT
                      *  *     *   * *             *          * *    **   **

Dnmt1mRNA           ATGCTCGTGAAGTCTGTCTAAAAGTTAAACGCATCT--ACCGTCCGGCCGACACCCATTT
Dnmt1amplicon       ATGCTCGTGAAGTCTGTCTAAAAGTTAAACGCATCT--ACCGTCCGGCCGACACCCATTT
Dnmt3amplicon       -TGGTCTTCTGCTGGAAAAGTGGATTCCACCTCTTT--TCTATCTTGCCAAAGGTCATCT
Dnmt2amplicon       ATGACCTCCTTTCCTAACAGTAGACTCTGTGACATTGAAATGCCCGTGCAAGAAAAAACG
                     **  *                   *         *       *    * *     *

Dnmt1mRNA           G-GGGCGTGACGCAGGCTTTCGCTCTGACTGGAATTTGGTCTACTGGTCAGATGAGATAC
Dnmt1amplicon       G-GGGCGTGACGCAGGCTTTCGCTCTGACTGGAATTTGGTCTACTGGTCAGATGAGATAC
Dnmt3amplicon       ACAAACCATCCATAATGATGTAAACAAATTGGAAAATGTTTGTTTGGCTTGTTGGGAAAA
Dnmt2amplicon       TTAGACCCCTATCTGGTGAAAGATATGTCTGATGAAGAATTGGCTCGTTATTTG---CTT
                         *                       **        *    * *     **

Dnmt1mRNA           ACAAACTGGAGTTAAACAAAGTTGTAGACAAGTGCGTCCTTGTCTGCAGCACTGCCATTG
Dnmt1amplicon       ACAAACTGGAGTTAAACAAAGTTGTAGACAAGTGCGTCCTTGTCTGCAGCACTGCCATTG
Dnmt3amplicon       AGGAACCTGTCCCCATCC--TTTTTTTGTAGGTGC-TTTGTGTCAACAGTGCAAACAAAG
Dnmt2amplicon       ACTGATAAAACATTAT-----TTAAGTATTGGCGCATTCTCGA-TGTACGACAAACATCC
                    *   *         *      **        * ** *    *     *   *   **

Dnmt1mRNA           ATGAACCCATTGAAGAATTTGTCCGGGGTGGACCTAACCGCATGTATTTCAATAAGGCC-
Dnmt1amplicon       ATGAACCCATTGAAGAATTTGTCCGGGGTGGACCTAACCGCATGTATTTCAATAAGGCC-
Dnmt3amplicon       ATTA--CTACGCACAATGTTTTCCAAAACCG--TTGATGGTTTTCATCTCTATTGCACCA
Dnmt2amplicon       GACACATCTTCCTGTTGTTTCACCAAAGCGTA-TACCCACTATGCAGAAGGAACTGGTT-
                       *              **  **                  *  *     *

Dnmt1mRNA           --TACAACCCGGCTGAGCGGGAATTTGAGCCGCCACCAGTGGAAGCCGAGCGAATCGGTT
Dnmt1amplicon       --TACAACCCGG------------------------------------------------
Dnmt3amplicon       TTTGTGGTTC--------------------------------------------------
Dnmt2amplicon       -CTGTACTTCAACACG--------------------------------------------
                      *      *
```

**Supplemental Figure 1** Alignment between the mRNA transcript sequence of DNMT 1 and the amplicon sequences used for cloning into the L4440 plasmid.

```
Dnmt2              TTCCATTTTCGTGAATTTATACTCTCTCCTGAAAGCATCAAAATACCCAATAGTCGAAGC
Dnmt2amplicon      ---------------------------------------------CCCAATAGTCGAAGC
Dnmt3amplicon      ------------------------------------------------GACAGCGACGCT
Dnmt1amplicon      -----------------------------------------------CACCCGACCCATT
                                                                       *

Dnmt2              AGATACTACCT-TGTAGCTAAAAA-GTGTACAGACTTCTCATTTGGATCTGAAAACGACA
Dnmt2amplicon      AGATACTACCT-TGTAGCTAAAAA-GTGTACAGACTTCTCATTTGGATCTGAAAACGACA
Dnmt3amplicon      ACTTATT--TT-TGTGGTTCGGAT-GACGATGATGGTGATGGAAATTTCGGAGTAGAAGC
Dnmt1amplicon      CTGTATTGGCTATGTCGTTGGTATTGCCTATAATGGCATCCTGCACAACAATTTGAATGC
                      ** *   * *** * *   *  *   *                  *

Dnmt2              TTATGACCTCCTTTCCTAACAGTAGACTCTGTGACATTGAAATGCCCGTGCAAGAAAAAA
Dnmt2amplicon      TTATGACCTCCTTTCCTAACAGTAGACTCTGTGACATTGAAATGCCCGTGCAAGAAAAAA
Dnmt3amplicon      TGATGGTCTTCTGCTGGAAAAGTGGATTCCACCTCTTTTCTAT---CTTGCCAAAGGTCA
Dnmt1amplicon      TCGTGAAGT-CTGTCTAAAAGTTAAACGCATCTACCGT--------CCGGCCGACACCCA
                   *  **   * **     **   *  *  *     *  *        *  **        *

Dnmt2              CGTTAGACCCCTATCTGGTGAAAGATATGTCTGATGAAGAATTGGCTCGTTATTTGCTTA
Dnmt2amplicon      CGTTAGACCCCTATCTGGTGAAAGATATGTCTGATGAAGAATTGGCTCGTTATTTGCTTA
Dnmt3amplicon      TCTACAAACCATCCATAATGATGTAAACAAAT--TGGAAAAT--GTTTGT--TTGGCTTG
Dnmt1amplicon      TTTGGGGCGTGACGCAGGCTTTCGCTCTGACT----GGAATTTGGTCTAC---TGGTCAG
                     *                            *       * *  *        * *

Dnmt2              CTGATAAAACATTATTTAAGTATTGGCGCATTCTCGAT--GTACGACAA-ACATCCGACA
Dnmt2amplicon      CTGATAAAACATTATTTAAGTATTGGCGCATTCTCGAT--GTACGACAA-ACATCCGACA
Dnmt3amplicon      TTGGGAAAAAGGAACCTGTCCCCATCCTTTTTTTGTAG--GTGCTTTGT-GTCAACAGTG
Dnmt1amplicon      ATGAGATACACAAACTGGAGTTAAACAAAGTTGTAGACAAGTGCGTCCTTGTCTGCAGCA
                    **  * *     *                ** *  *   ** *           *

Dnmt2              CATCTTCCTGTTGTTTCACCAAAGCGTATACCCACTATGCAGAAGGAACTGGTTCTGTAC
Dnmt2amplicon      CATCTTCCTGTTGTTTCACCAAAGCGTATACCCACTATGCAGAAGGAACTGGTTCTGTAC
Dnmt3amplicon      CAAACAAAGATTACTACGC-ACAATGTTTTCCAAAACCGTTGATGGTTTTCAT-CTCTAT
Dnmt1amplicon      CTGCCATTGATGAACCCATTGAAGAATTTGTCCGGGGTG-GACCTAACCGCATGTATTTC
                   *         *     *     *   * *  *      *             *    *

Dnmt2              TTCAACACGACCCGAACGAGCCATTTCACCAAAAATTCGCTGAATTTAAAGAAGATGAAG
Dnmt2amplicon      TTCAACACG---------------------------------------------------
Dnmt3amplicon      TGCACCATTTGTGGTTC-------------------------------------------
Dnmt1amplicon      AATAAGGCCTACAACCCGG-----------------------------------------
                      *
```

**Supplemental Figure 2** Alignment between the mRNA transcript sequence of DNMT 2 and the amplicon sequences used for cloning into the L4440 plasmid.

```
Dnmt3_mRNA          ATTCAGACGATTGTTATGACAGCGACGCTACTTATT--TT-TGTGGTTCGGAT-GACGAT
Dnmt3amplicon       -----------------GACAGCGACGCTACTTATT--TT-TGTGGTTCGGAT-GACGAT
Dnmt2amplicon       --------------CCCAATAGTCGAAGCAGATACTACCT-TGTAGCTAAAAA-GTGTAC
Dnmt1amplicon       ----------------CACCCGACCCATTCTGTATTGGCTATGTCGTTGGTATTGCCTAT
                                         *          ** *   * *** * *   *  *   *

Dnmt3_mRNA          GATGGTGATGGAAATTTCGGAGTAGAAGCTGATGGTCTTCTGCTGGAAAAGTGGATTCCA
Dnmt3amplicon       GATGGTGATGGAAATTTCGGAGTAGAAGCTGATGGTCTTCTGCTGGAAAAGTGGATTCCA
Dnmt2amplicon       AGACTTCTCATTTGGATCTGAAAACGACATTATGACCTCCTTTCCTAACAGTAGACTCTG
Dnmt1amplicon       AATGGCATCCTGCACAACAATTTGAATGCTCGTGAAGT-CTGTCTAAAAGTTAAACGCAT
                                     *           *  **   * **     **   *  *  *

Dnmt3_mRNA          CCTCTTTTCTAT---CTTGCCAAAGGTCATCTACAAACCATCCATAATGATGTAAACAAA
Dnmt3amplicon       CCTCTTTTCTAT---CTTGCCAAAGGTCATCTACAAACCATCCATAATGATGTAAACAAA
Dnmt2amplicon       TGACATTGAAATGCCCGTGCAAGAAAAAACGTTAGACCCCTATCTGGTGAAAGATATGTC
Dnmt1amplicon       CTACCGT--------CCGGCCGACACCCATTTGGGGCGTGACGCAGGCTTTCGCTCTGAC
                       *  *        *  **        *  *

Dnmt3_mRNA          TTGGAAAATGTTTGTTTG--GCTTGTTGGGAAAAAGGAACCTGTCCCCATCCT--TTTTT
Dnmt3amplicon       TTGGAAAATGTTTGTTTG--GCTTGTTGGGAAAAAGGAACCTGTCCCCATCCT--TTTTT
Dnmt2amplicon       TGATGAAGAATTGGCTCGTTATTTGCTTACTGATAAAA---------CATTAT--TTAAG
Dnmt1amplicon       TGGAATTTGGTCTACTGG---TCAGATGAGATACACAAACTGGAGTTAAACAAAGTTGTA
                    *         *    * *      * *     * *  *          *      **

Dnmt3_mRNA          TGTAGGTGCTTTGT-GTCAACAGTGCAAACAAAGATTAC-TACGCACAATGTTTTCCAAA
Dnmt3amplicon       TGTAGGTGCTTTGT-GTCAACAGTGCAAACAAAGATTAC-TACGCACAATGTTTTCCAAA
Dnmt2amplicon       TATTGGCGCATTCT-CGATGTACGACAAACATCCGACACATCTTCCTGTTGTTTCACCAA
Dnmt1amplicon       GACAAGTGCGTCCTTGTCTGCAGCACTGCCATTGATGAACCCATTGAAGAATTTGTCCGG
                         * ** *  *       *   *   **      *             ***  *

Dnmt3_mRNA          ACCGTTGATGGTTTTCATCTCTATTGCACCA--TTTGTGGTTCAAAATCAAATGCCTGTG
Dnmt3amplicon       ACCGTTGATGGTTTTCATCTCTATTGCACCA--TTTGTGGTTC-----------------
Dnmt2amplicon       AGCGT--ATACCCACTATGCAGAAGGAACTGGTTCTGTACTTCAACACG-----------
Dnmt1amplicon       GGTGGACCTAACCG-CATGTAT-TTCAATAAGGCCTACAACCCGG---------------
                       *    *       **         *       *      *
```

**Supplemental Figure 3** Alignment between the mRNA transcript sequence of DNMT 3 and the amplicon sequences used for cloning into the L4440 plasmid.

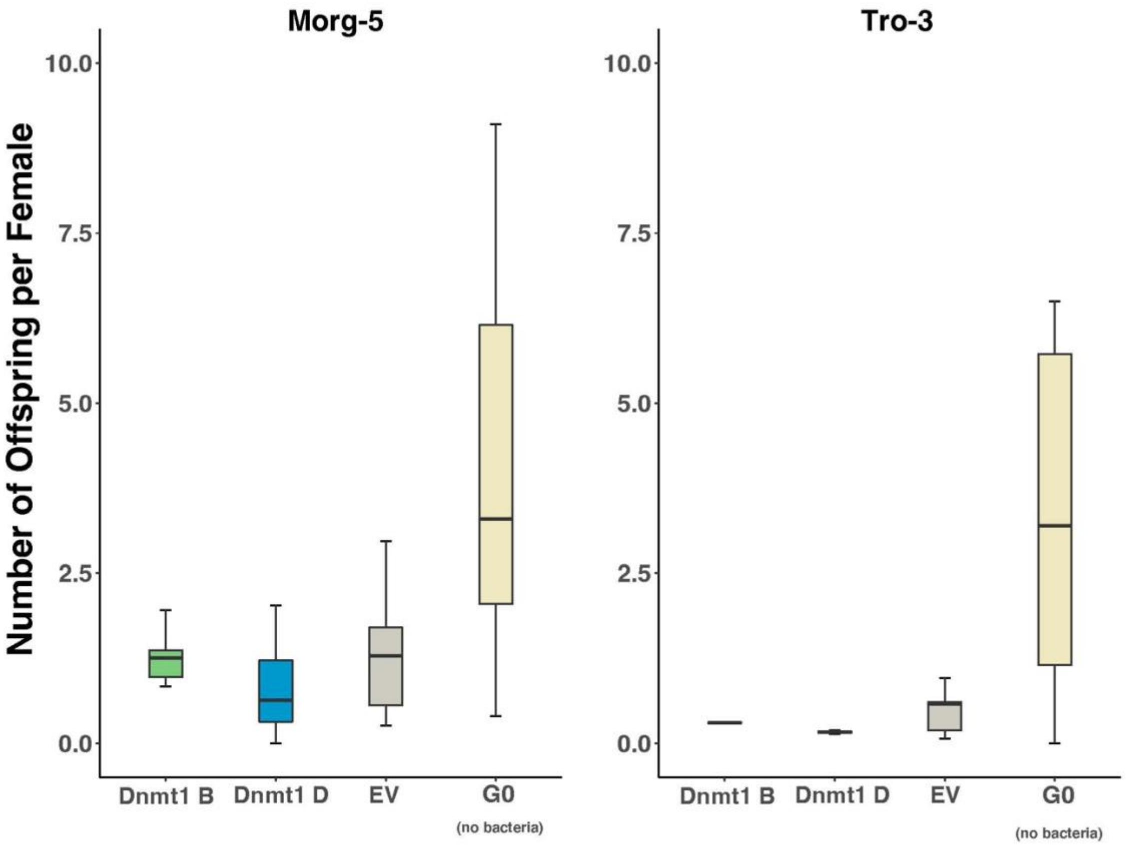


**Supplemental Figure 4** Standardized boxplots of the number of neonates produced each day, total number of neonates for each treatment divided by the number of surviving females. Animals treated with the DNMT1 B vector are shown in green, DNMT1 D vector in blue, and EV in gray. No significant difference was detected between vector treatments. When compared to the maternal generation, $G_0$, all three vector treatments in both clones produced significantly fewer offspring over a similar timeframe.

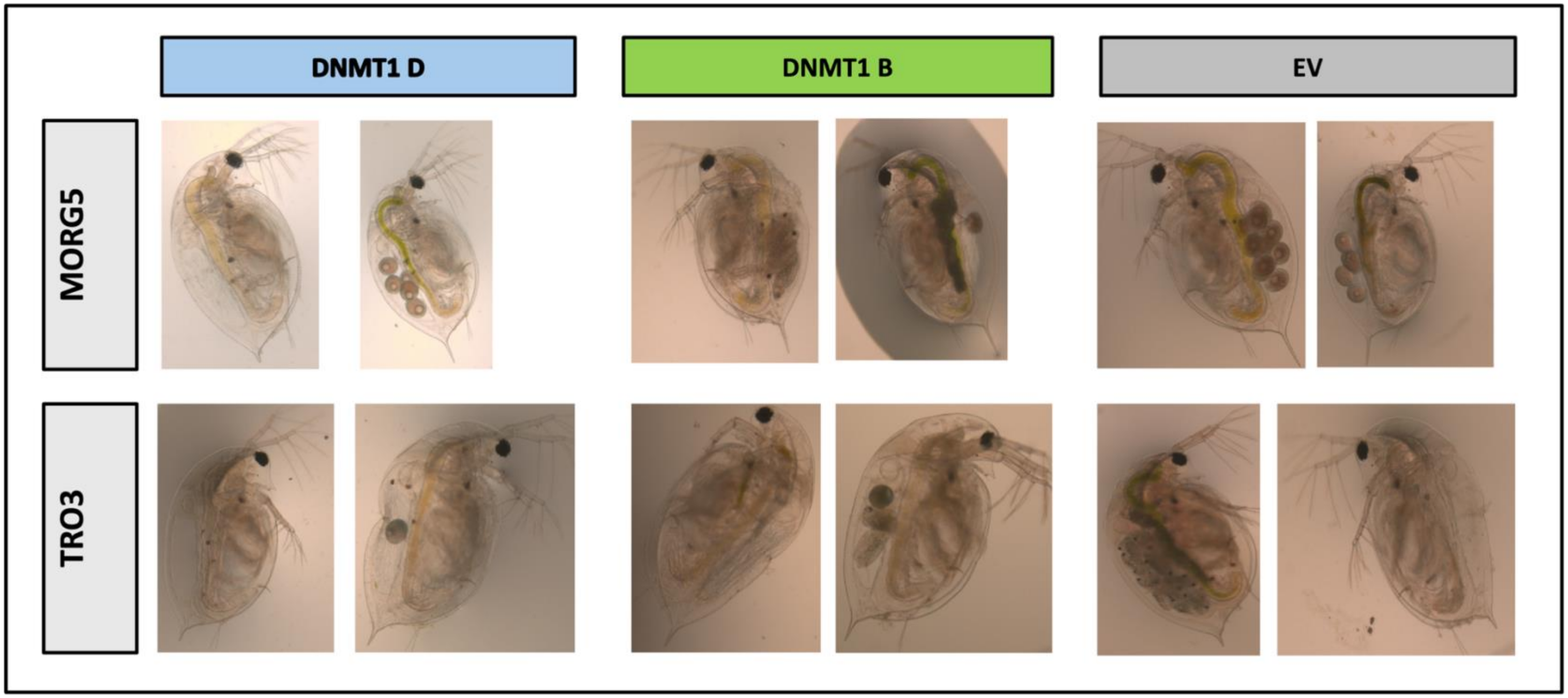


**Supplemental Figure 5** Photographs taken on day 10 of the RNAi bacterial feeding regime. TRO3 animals show more obvious signs of stress: loss of pigmentation in the carapace, dark yellow to brown coloring in the gut, decreased egg production, and a deformity in the carapace. MORG5 animals do not show the same signs of distress in any treatment. Vitellogenesis and egg production continued unimpaired, gut coloring indicated constant filter feeding, and no physical deformities were observed.

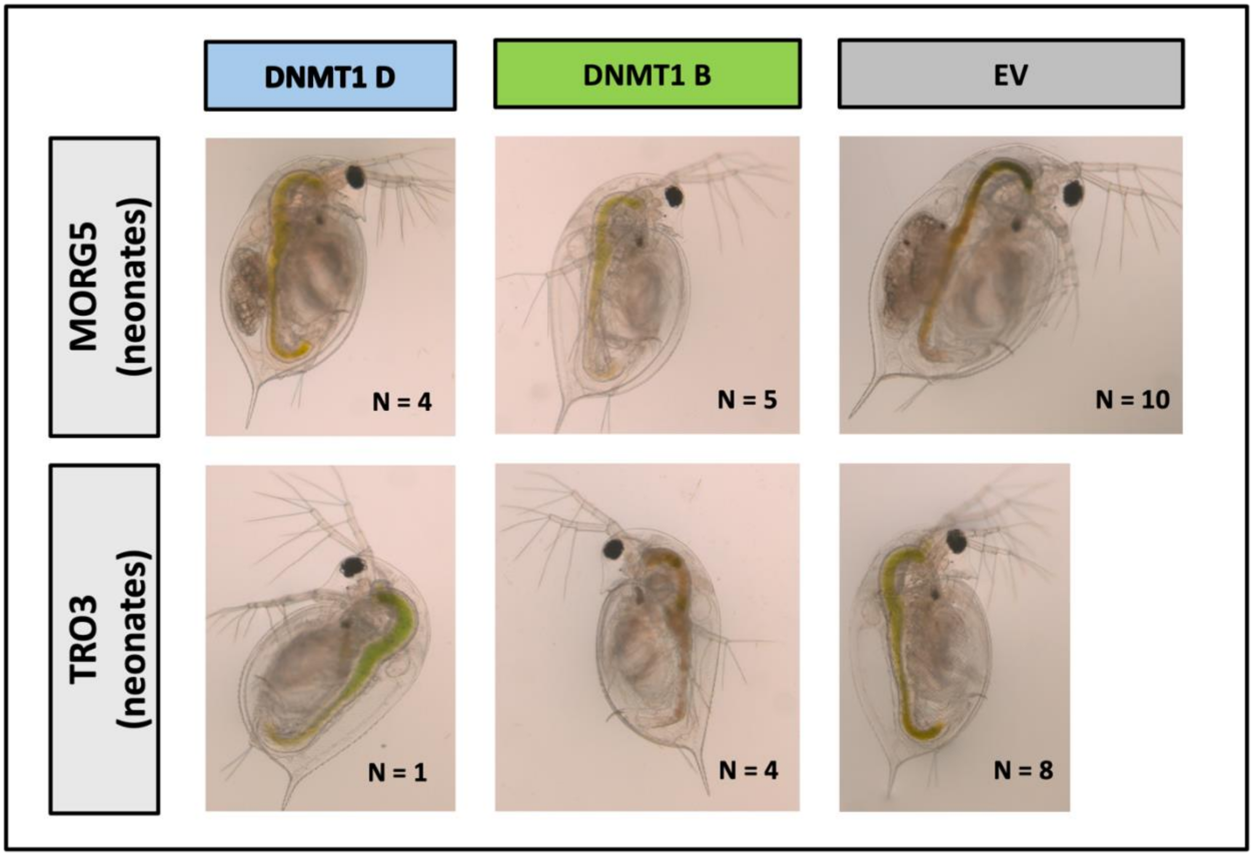


**Supplemental Figure 6** Neonates born into during the bacterial feeding regime were collected (n = 10 per treatment per clone) and treated with the same vector as their mother. The N represents how many of the original cohort survived by day 12 when the photographs were taken. Developmentally, all surviving offspring appear to have defined brood chambers but only MORG5, DNMT1 D, and EV produced eggs. Maturation was delayed in both treatments by approximately 2.5 days.

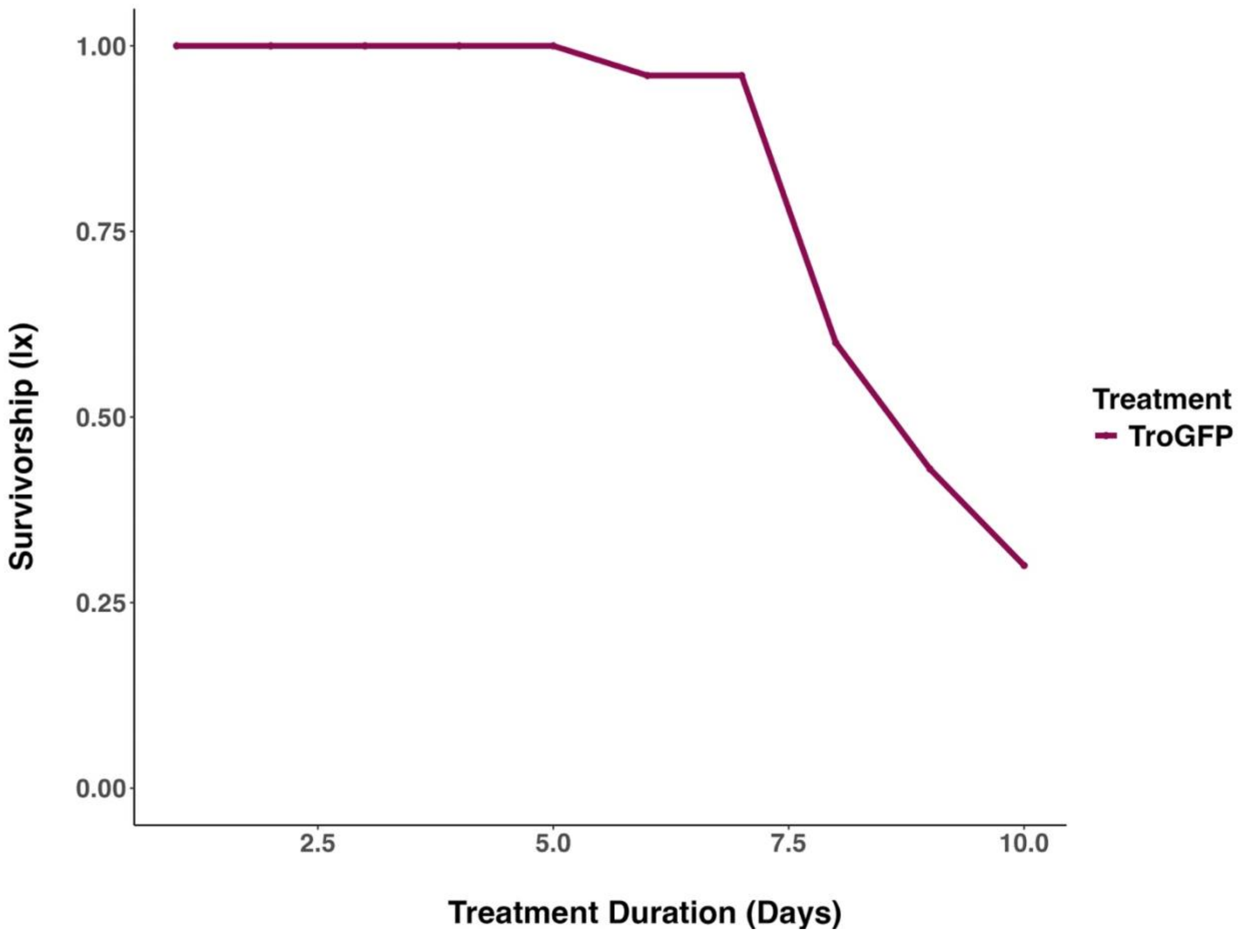


**Supplemental Figure 7** Plot showing the effects of the GFP treatment on survivorship (lx) of TRO3 animals. TRO3 GFP survivorship was significantly reduced ($\chi^2$ = 14.046, $p < 0.029$, $df = 6$) when compared with animals from Experiment 2.

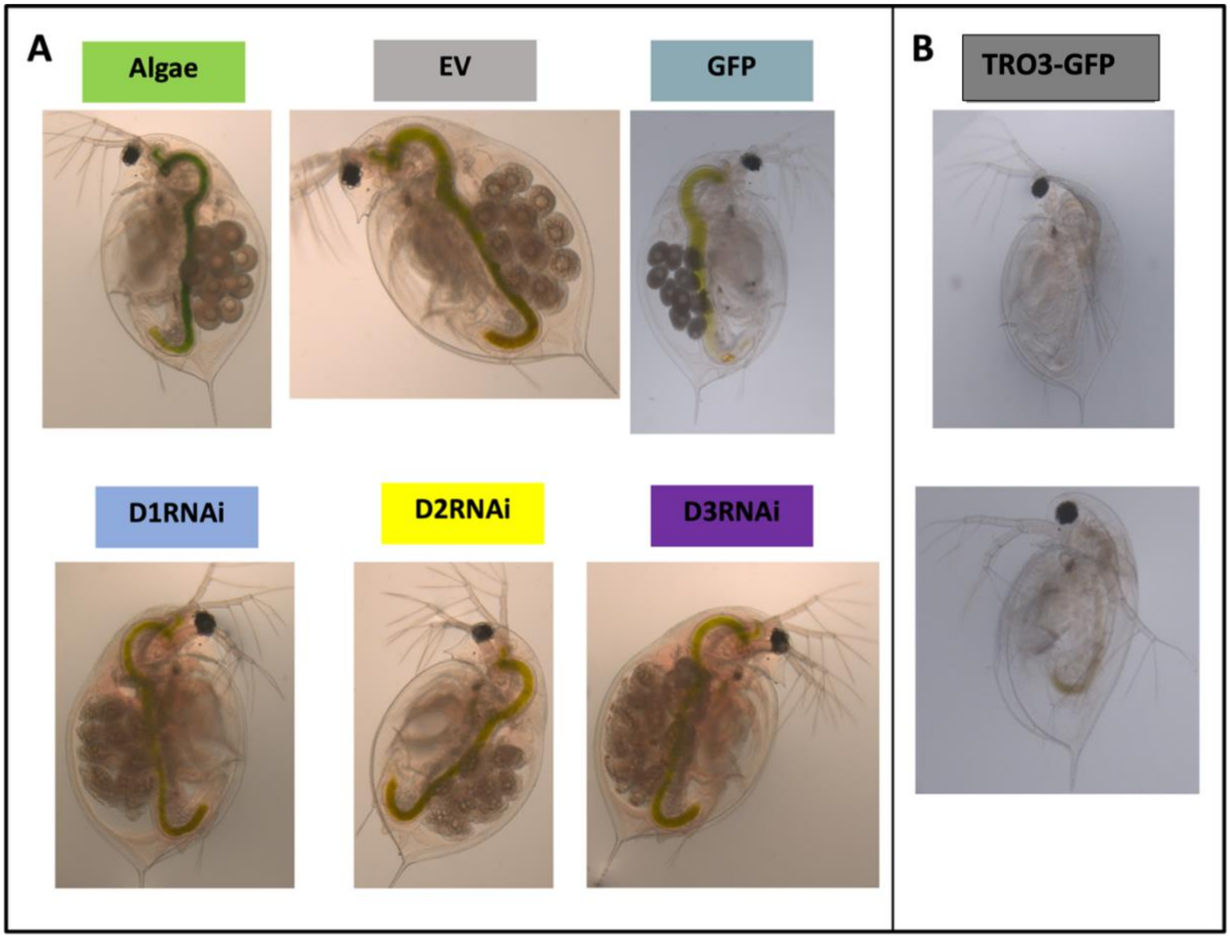


**Supplemental Figure 8** Photographs taken on day 10 of the RNAi bacterial feeding regime. **A)** Photographs of the MORG5 clone from each treatment Algae (green), Empty Vector (grey), L4440 with GFP (turquoise), and each DNA methyltransferase RNAi treatment 1 (blue), 2 (yellow), and 3 (purple). On days 2 and 3, animals from all three DNMT RNAi treatments experienced two ecdysis events, both premature and without the release of neonates. Gut coloring for all six treatments is light to bright green, indicating constant filter feeding of fresh algae. Vitellogenesis and egg production continued unimpaired with normal egg development. All three DNMT RNAi treatments resulted in animals producing high levels of hemoglobin, giving them a pink coloring. **B)** Photographs of surviving TRO3 animals from the GFP plasmid vector treatment. Animals appear translucent; gut coloring is either dark yellow-brown or clear, indicating inconsistent, minimal filtering or a complete halt to filter feeding. Despite reaching maturation at the start of the experiment, no offspring were produced, and vitellogenesis was not observed during the 10-day feeding regime.

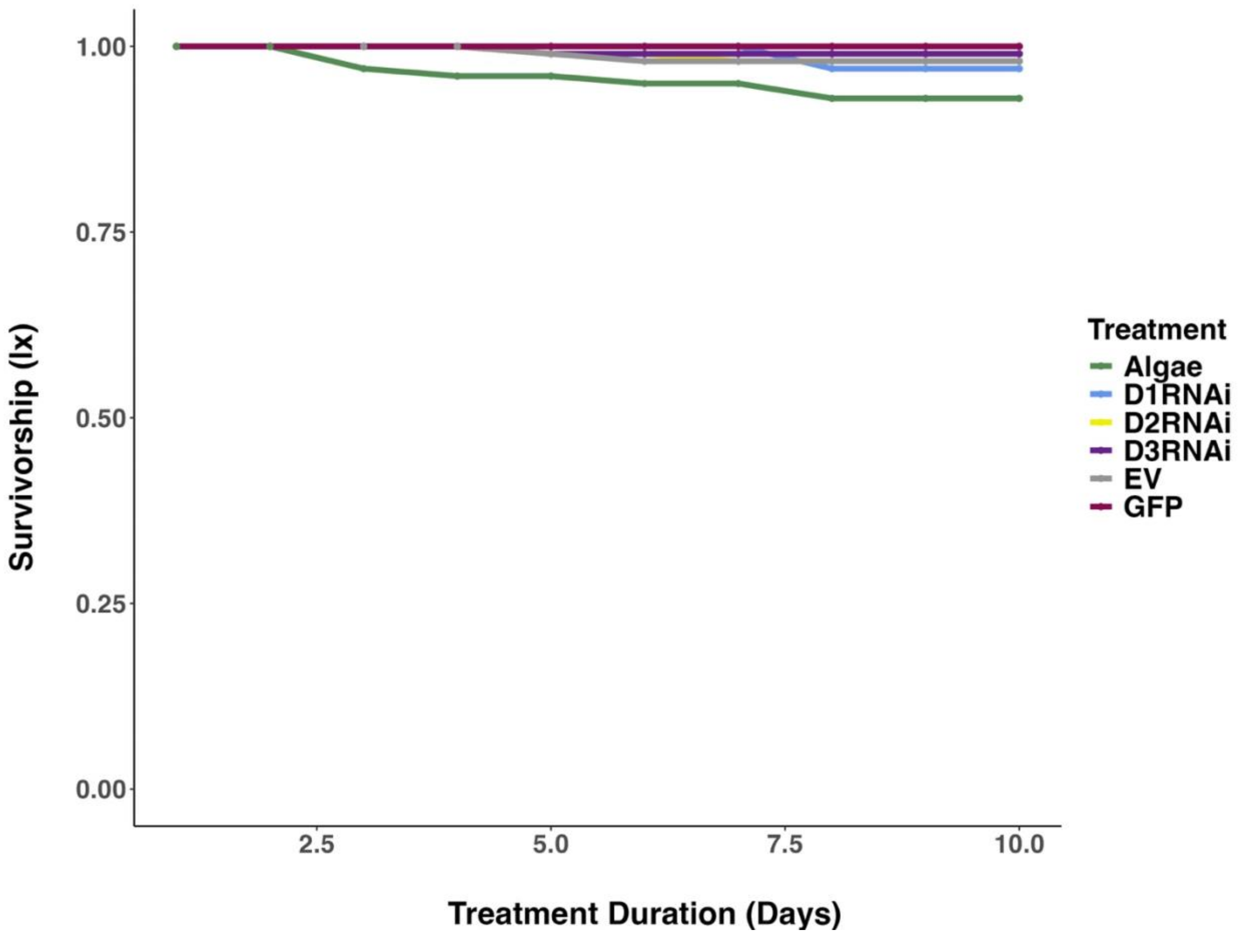


**Supplemental Figure 9** Plot showing the effects of each treatment on survivorship (*lx*) for both clones, MORG5 and TRO3. Treatment identification is as follows: Algae (green), Empty Vector (grey), L4440 with GFP (pink), and each DNA methyltransferase RNAi treatment 1 (blue), 2 (yellow), and 3 (purple). The was no significant effect of treatment on survivorship observed for the MORG5 clone ($\chi^2$ = 0.366, p = 0.996, *df* = 5).

**Supplemental Table 1: DNMT1 Dunn's Test Multiple Comparison**

Kruskal-Wallis: $\chi^2 = 28.99$, $df = 5$, $p < 0.0001$

| | Algae | D1RNAi | D2RNAi | D3RNAi | EV |
|---|---|---|---|---|---|
| D1RNAi | -2.334<br>**0.042** | | | | |
| D2RNAi | -2.453<br>**0.053** | -0.119<br>0.905 | | | |
| D3RNAi | -3.089<br>**0.01** | -0.745<br>0.563 | -0.636<br>0.562 | | |
| EV | -0.634<br>0.606 | 1.698<br>0.134 | 1.817<br>0.115 | 2.453<br>**0.043** | |
| GFP | -4.717<br>**0.000** | -2.383<br>**0.043** | -2.264<br>**0.044** | -1.628<br>0.141 | -4.081<br>**0.000** |

Dunn's pairwise *z* statistic and p-value, significant p-values are in bold
Alpha = 0.05
Method = Benjamini – Hochberg

**Supplemental Table 2: DNMT2 Dunn's Test Multiple Comparison**
Kruskal-Wallis: $\chi^2 = 23.27$, $df = 5$, $p < 0.0002$

| | Algae | D1RNAi | D2RNAi | D3RNAi | EV |
|---|---|---|---|---|---|
| D1RNAi | -4.004<br>**0.000** | | | | |
| D2RNAi | -2.516<br>**0.035** | 1.489<br>0.256 | | | |
| D3RNAi | -3.012<br>**0.013** | 0.992<br>0.438 | -0.496<br>0.775 | | |
| EV | -2.600<br>**0.035** | 1.405<br>0.267 | -0.084<br>0.933 | 0.412<br>0.785 | |
| GFP | -4.283<br>**0.000** | -0.280<br>0.836 | -1.768<br>0.193 | -1.272<br>0.305 | -1.684<br>0.197 |

Dunn's pairwise *z* statistic and p-value, significant p-values are in bold
Alpha = 0.05
Method = Benjamini – Hochberg

**Supplemental Table 3: DNMT3 Dunn's Test Multiple Comparison**
Kruskal-Wallis: $\chi^2 = 34.77$, $df = 5$, $p < 0.0001$

| | Algae | D1RNAi | D2RNAi | D3RNAi | EV |
|---|---|---|---|---|---|
| D1RNAi | -2.921<br>**0.011** | | | | |
| D2RNAi | -2.348<br>**0.035** | 0.573<br>0.708 | | | |
| D3RNAi | -2.719<br>**0.014** | 0.203<br>0.839 | -0.370<br>0.821 | | |
| EV | -2.055<br>**0.067** | 0.867<br>0.579 | 0.294<br>0.824 | 0.664<br>0.691 | |
| GFP | -5.787<br>**0.000** | -2.865<br>**0.010** | -3.439<br>**0.003** | -3.068<br>**0.008** | -3.732<br>**0.001** |

Dunn's pairwise $z$ statistic and p-value, significant p-values are in bold
Alpha = 0.05
Method = Benjamini – Hochberg
